# Genome analysis of the steroid-degrading denitrifying *Denitratisoma* oestradiolicum DSM 16959 and *Denitratisoma* sp. strain DHT3

**DOI:** 10.1101/710707

**Authors:** Yi-Lung Chen, Sean Ting-Shyang Wei, Yin-Ru Chiang

## Abstract

Steroid hormones (androgens and estrogens) are crucial for development, reproduction, and communication of multicellular eukaryotes. The ubiquitous distribution and persistence of steroid hormones in our ecosystems have become an environmental issue due to the adverse effects on wildlife and humans upon long-term exposure. Microbial degradation is critical for the removal of steroid hormones from ecosystems. The aerobic degradation pathways for androgens and estrogens and the anaerobic degradation pathway for androgen have been studied into some details; however, the mechanism for anaerobic estrogen degradation remains completely unknown. Here, we presented the circular genomes of *D. oestradiolicum* DSM 16959 and *Denitratisoma* sp. strain DHT3, two betaproteobacteria capable of anaerobic estrogen degradation. We identified the genes involved in steroid transformation and in the anaerobic 2,3-*seco* pathway in both genomes. Additionally, the comparative genomic analysis revealed that genes exclusively represented in estrogen-degrading anaerobes might play a role in anaerobic estrogen catabolism.

## Introduction

Steroid sex hormones, including androgens and estrogens, play essential roles in the physiology, development, reproduction, and behaviors of vertebrates. The occurrence and persistence of steroid sex hormones in our environments, especially in aquatic ecosystems, result in interruption for animal physiology and behavior. Lambert et al (2015) showed that for amphibian, long-term exposure to estrogens even at extremely low concentration lead to a female-dominated frog population. Moreover, estrogens not only act as endocrine disruptor but have also been classified as Group 1 carcinogens by the World Health Organization (http://monographs.iarc.fr/ENG/Classification/latest_classif.php).

The ability to produce steroid sex hormones is only conserved in eukaryotes, but interestingly, bacteria appear to be the major steroid degraders in the biosphere (Holert et al., 2018), and adopt various catabolic pathways to degrade these recalcitrant compounds depending on the oxygen availability (Chen *et al*., 2017; Casabon *et al*., 2017). In general, under aerobic condition, bacteria adopt the 9,10-seco pathway (Bergstrand *et al*., 2016) and the 4,5-seco pathway (Chen *et al*., 2017, 2018) to degrade androgens and estrogens, respectively; under anaerobic condition, denitrifying bacteria degrade androgens through the 2,3-seco pathway (Wang *et al*., 2013; Yang *et al*., 2016). To date, only betaproteobacterial *Denitratisoma oestradiolicum* DSM 16959 (Fahrbach *et al*., 2006) and gammaproteobacterial *Steroidobacter denitrificans* DSM 18526 are capable of anaerobic estrogen degradation (Fahrbach *et al*., 2008); however, their anaerobic catabolic mechanism remains unclear. In this study, we sequenced and annotated the genome of two estrogen-degrading anaerobes–D. *oestradiolicum* DSM 16959 and *Denitratisoma* sp. strain DHT3–from a municipal wastewater treatment plant. The comparative genomic analysis showed that these two betaproteobacteria harbor genes involved in anaerobic degradation for steroidal ABCD rings. Moreover, the genes only identified in three estrogen-degrading anaerobes might play important roles in estrogens catabolism.

## Material and Methods

### Genome sequencing, assembling and annotation

Genomic DNA of strain DSM 16959 and strain DHT3 were extracted using the Easy Tissue & Cell Genomic DNA Purification Kit (GeneMark, Taiwan). The workflow for genome sequencing and bioinfomatic analysis is availabe in Figure 1. For strain DHT3, purified genomic DNA was sequenced on two platforms: Illumina HiSeq 2500 (Illumina Inc., San Diego, CA, USA) and PacBio RSII (Pacific Biosciences, CA, USA). Two Illumina TruSeq^®^ DNA PCR-Free libraries with fragment size around 170 bp (fragment library) and 555 bp (jumping library) were prepared for Illumina paired-end sequencing (2x 125bp). Subsequently, under the default settings, the adapter sequences were removed using cutadapt (v1.4.2; Martin, 2011) and then low-quality bases were trimmed by Seqtk (v1.2-r94; https://github.com/lh3/seqtk). After these two trimming steps, the sequences longer than 35 nucleotides were included for succeeding analysis. For Pacific Biosciences (PacBio) platform, one SMRT cell was used for sequencing. Reads acquired from PacBio were assembled *de novo* using RS_HGAP_assembly.3 protocol included in SMART Portal (version 2.3.0). These assembled contigs were further corrected by the Illumina reads using bowtie2 (version 2.2.3; Langmead and Salzberg, 2012) for alignment, and Samtools (Version: 0.1.19-44428cd; Li *et al*., 2009) and bcftools (https://github.com/samtools/bcftools) to extract consensus sequence with default setting. The genome of the strain DSM 16959 was only sequenced on Illumina HiSeq 2500 platform. Two Illumina TruSeq^®^ DNA PCR-Free libraries (fragment library and jumping library) were also prepared. Same bioinformatic analysis processes were applied as to strain DHT3 except for adopting AllPaths-LG (Gnerre *et al*., 2011) on strain DHT3 genome assemble.

**Figure 1.**
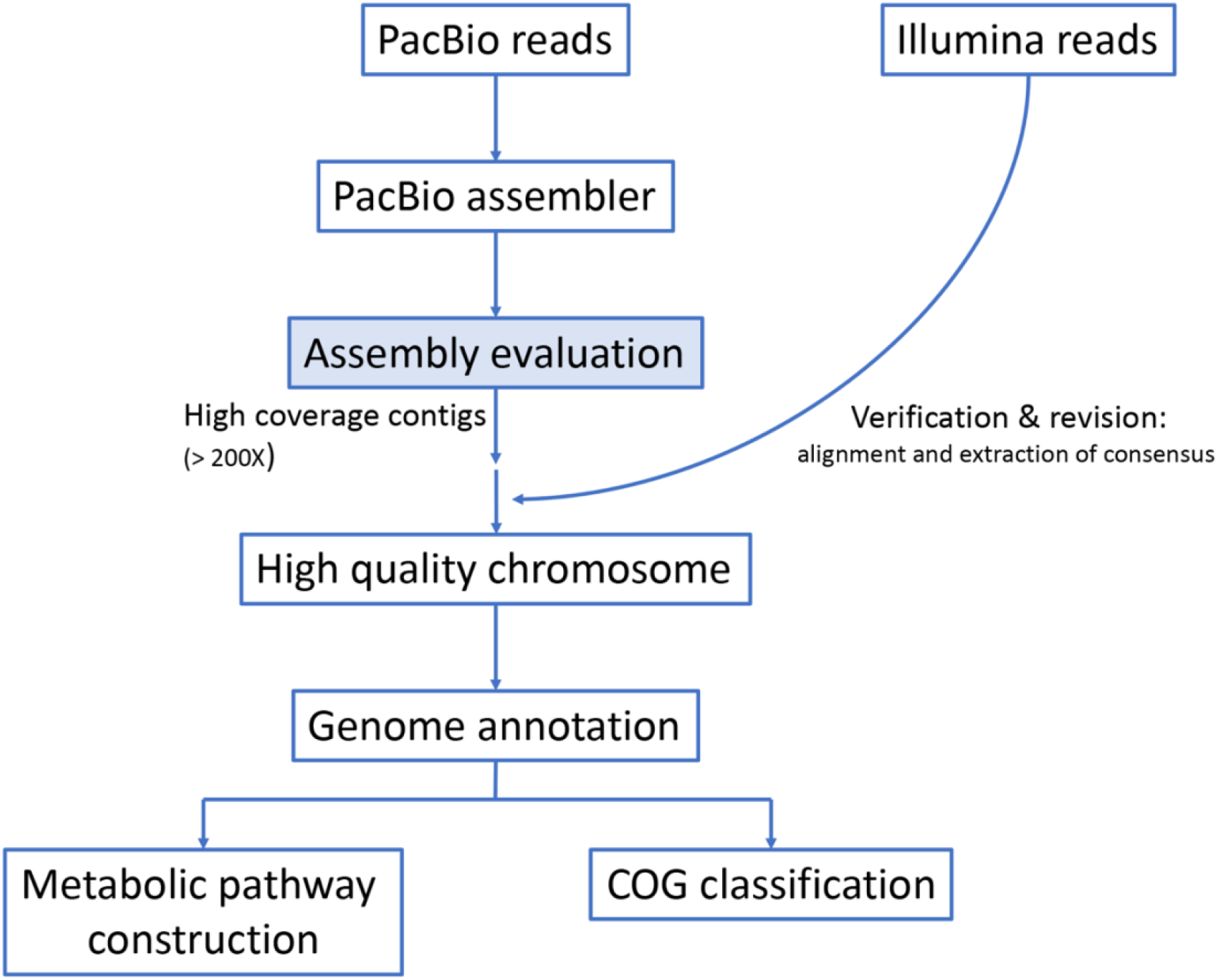
**Genome sequencing and** bioinfomatic workflow for the strain DHT3 and its functional annotation.

The whole genomes were annotated using the NCBI Prokaryotic Genome Annotation Pipeline (Tatusova *et al*., 2016) and the protein-coding genes were classified into COG category by the eggNOG-mapper (Huerta-Cepas *et al*., 2017). For constructing the metabolic pathway *in silico*, the KEGG BlastKOALA was applied (Kanehisa *et al*., 2016).

### Comparative genomic analysis

The bacterial comparative genomic analysis was based on the gene orthologue shared between the genomes of 6 steroid-degrading aerobes and anaerobes, including *Sphingomonas* sp. strain KC8 (Chen *et al*., 2017), *Sterolibacterium denitrificans* DSM 13999 (Warnke *et al*., 2017), *Thauera terpenica* strain 58Eu (Foss and Harder, 1998), strain DHT3 (this study), *Denitratisoma oestradiolicum* DSM 16959 (this study) and *Steroidobacter denitrificans* DSM 18526 (Yang *et al*., 2016). Their phylogeny and physiological features on steroid degradation, as well as genome accession numbers are summarized in the Table 1. Their total protein sequences of coding region were uploaded to the web server, OrthoVenn2 (Xu *et al*., 2019) for comparing and annotating the gene content based on their orthology under the parameters: e-value: 1e-15 and inflation value: 1.5.

**Table 1.** Steroid degradation capacity under aerobic and anaerobic conditions of six different steroid degraders. +: growth, −: no growth, ND: not determined.

| Steroid | Alpha-proteobacteria |  | Beta-proteobacteria |  |  |  |  |  |  |  | Gamma-proteobacteria |  |
| --- | --- | --- | --- | --- | --- | --- | --- | --- | --- | --- | --- | --- |
|  | *Sphingomonas* sp.<br>strain KC8* |  | *Denitratisoma* sp.<br>strain DHT3 |  | *Denitratisoma*<br>*oestradiolicum*<br>DSM 16959 |  | *Sterolibacterium*<br>*denitrificans* DSM<br>13999‡ |  | *Thauera terpenica*<br>strain 58Eu§ |  | *Steroidobacter*<br>*denitrificans* DSM<br>18526\|\| |  |
|  | CP016306¶ |  | CP029914¶ |  | NCXS000000000¶ |  | LT837803~LT837806¶ |  | NZ_ATJV01000001¶ |  | NZ_CP011971¶ |  |
|  | aerobic | anaerobic | aerobic | anaerobic | aerobic | anaerobic | aerobic | anaerobic | aerobic | anaerobic | aerobic | anaerobic |
| Cholesterol | ND | - | ND | - | ND | - | + | ND | ND | ND | ND | - |
| 17β-Estradiol | + | - | ND | + | - | + | ND | ND | ND | ND | - | + |
| Estrone | + | - | ND | ND | - | + | ND | ND | ND | ND | - | + |
| Testosterone | + | - | ND | ND | - | - | ND | + | ND | + | + | + |
| 4-androstene-3,17-dione | ND | - | ND | ND | - | - | ND | ND | ND | ND | + | + |
| 1,4-androstene-3,17-dione | ND | - | ND | ND | ND | ND | ND | ND | ND | ND | ND | ND |

\*Strain KC8 is an obligate aerobe and its steroid degradation ability can be referred to (Roh and Chu, 2010).
‡DSM 1399 is a facultative anaerobe and its steroid degradation ability can be referred to (Fahrbach *et al.*, 2006; Tarlera and Denner, 2003)(Wang *et al.*, 2014)
§Strain 58Eu is a facultative anaerobe and its steroid degradation ability can be referred to (Yang *et al.*, 2016).
|| DSM 18526 is a facultative anaerobe and its steroid degradation ability can be referred to (Yang *et al.*, 2016)(Fahrbach *et al.*, 2008).
¶ Accession number in the GenBank of NCBI.

## Results and Discussion

The phylogenetic analysis showed that strain DHT3 displayed highest 16S rRNA gene similarity (97.5 %) to D. *oestradiolicum* DSM 16959 (Fahrbach *et al*., 2006), suggesting that this strain belongs to the genus *Denitratisoma.* Therefore, this microorganism is named as *Denitratisoma* sp. strain DHT3 in this study.

### Genome of *Denitratisoma* sp. strain DHT3

In the present study, we obtain the high-quality circular genome of the strain DHT3, which was sequenced by two sequencing technologies. The Illumina and PacBio sequencing systems generated 11,617,819 reads (read length 125×2 bp) and 68,673 reads (read length 15,254-bp), respectively. After the quality trimming, the total read length from high-throughput sequencing was ~2,503 Mbp. Through PacBio sequencing, 19 contigs with N_50_ as 3.7 Mbps were obtained. After correction by the Illumina reads, one chromosome with 3.7 Mbps was revealed with 223-fold coverage of the genome (Table 2). This genome sequence is available in NCBI with the accession number of CP020914.

**Table 2.**
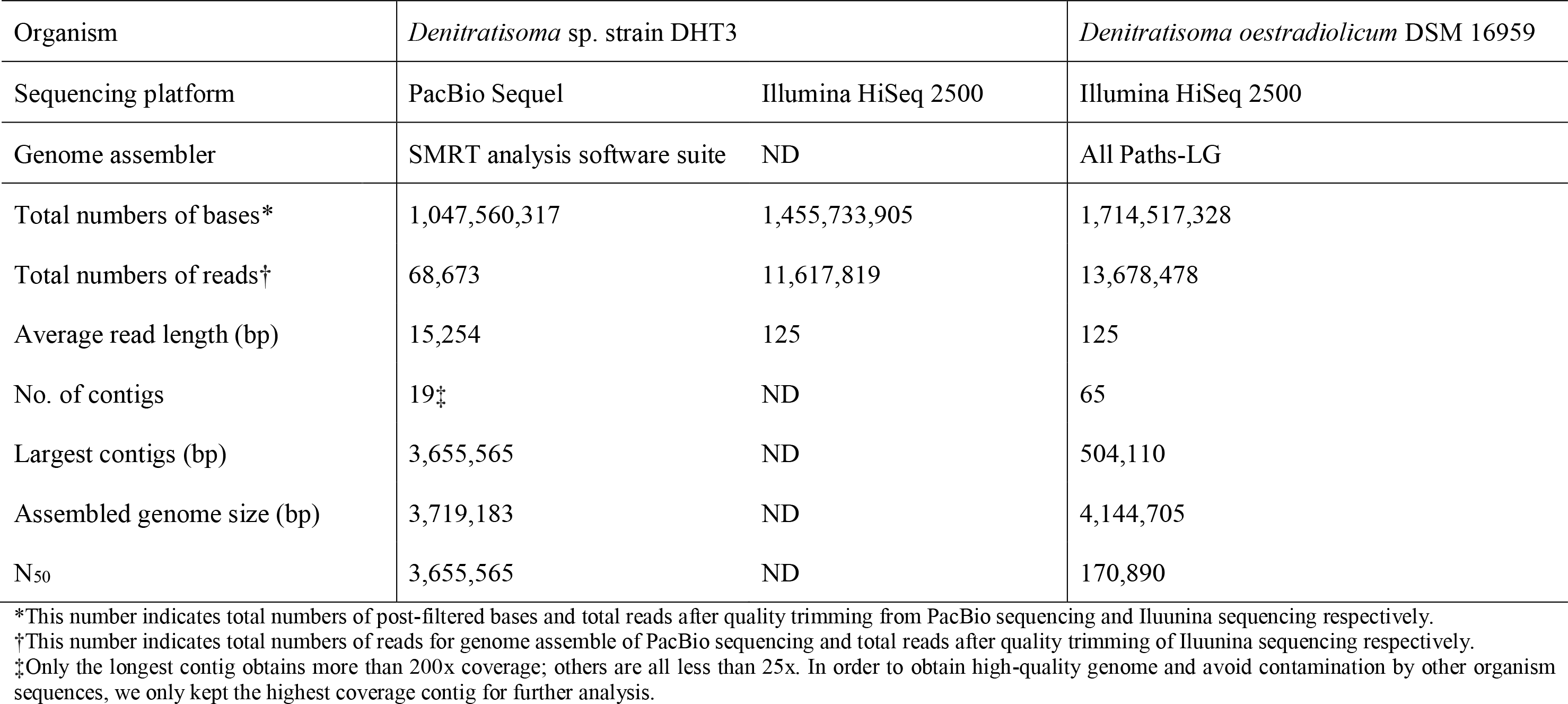
Summary of the quality trimming and assembly results. ND: not determined.

| Organism | *Denitratisoma* sp. strain DHT3 |  | *Denitratisoma oestradiolicum* DSM 16959 |
| --- | --- | --- | --- |
| Sequencing platform | PacBio Sequel | Illumina HiSeq 2500 | Illumina HiSeq 2500 |
| Genome assembler | SMRT analysis software suite | ND | All Paths-LG |
| Total numbers of bases* | 1,047,560,317 | 1,455,733,905 | 1,714,517,328 |
| Total numbers of reads† | 68,673 | 11,617,819 | 13,678,478 |
| Average read length (bp) | 15,254 | 125 | 125 |
| No. of contigs | 19‡ | ND | 65 |
| Largest contigs (bp) | 3,655,565 | ND | 504,110 |
| Assembled genome size (bp) | 3,719,183 | ND | 4,144,705 |
| N<sub>50</sub> | 3,655,565 | ND | 170,890 |

\*This number indicates total numbers of post-filtered bases and total reads after quality trimming from PacBio sequencing and Illumina sequencing respectively.
†This number indicates total numbers of reads for genome assemble of PacBio sequencing and total reads after quality trimming of Illumina sequencing respectively.
‡Only the longest contig obtains more than 200x coverage; others are all less than 25x. In order to obtain high-quality genome and avoid contamination by other organism sequences, we only kept the highest coverage contig for further analysis.

Based on NCBI annotation service, strain DHT3 chromosome is 3,655,661 bp with a G+C content of 64.9 %. Up to 3,246 protein coding genes, three rRNA operons (5S, 16S and 23S), 51 tRNA for all 21 amino acids, including selenocysteine, 1 CRISPR array and 71 pseudogenes are identified. Through eggNOG-mapper analysis, 2,917 protein-coding genes are classified into COG categories and the code S (function unknown) is the most abundant group (644 genes). This result implies the physiological traits and gene functions of the DHT3 are yet to be fully explored.

Bacterial steroid degradation requires coenzyme A (CoA) for the activation of the recalcitrant structures through β-oxidation reactions (Warnke *et al*., 2017; Casabon *et al*., 2017; Wu et al., 2019). We identified several genes involved in this β-oxidation (B9N43_01490 ~ 01520, 03830 and 04285).

In the strain DHT3 genome, we noted the gene cluster (B9N43_4420~4465) for 3-(7a-methyl-1,5-dioxooctahydro-1H-inden-4-yl)propanoic acid (HIP) catabolism (namely the steroid C/D-rings degradation) We also identified the genes involved in the steroid A/B-rings degradation through the 2,3-seco pathway, including the gene cluster encoding the 1-testosterone hydratase/dehydrogenase (B9N43_01910~1920) as well as the steroid dehydrogenase genes involved in steroid A-ring degradation [e.g., B9N43_03425 and _11350 (encoding putative 3α-hydroxysteroid dehydrogenase); B9N43_04410 (encoding putative 3-oxosteroid Δ^1^-dehydrogenase); B9N43 _16370 (coding for putative 3β-hydroxysteroid dehydrogenase); and B9N43_15155 (encoding putative 3-oxosteroid Δ^4^-dehydrogenase)]. Moreover, 4 sets of gene clusters encoding the putative steroid C25 dehydrogenase (B9N43_01670~01680; _5455~5470; _11160~11175; and _15060~15070) were identified. This molybdoeznyme is known to mediate anaerobic hydroxylation reactions on the tertiary carbons of the side chain of various sterols (Warnke et al., 2017) but is not reportedly involved in the steroidal core-ring degradation. The functions of these steroid C25 dehydrogenase genes remain further investigation.

Surprisingly, the DHT3 genome lacks the complete gene sets for glycolysis and TCA cycle (6-phophofructokinas and isocitrate dehydrogenase are absent for this central carbohydrate metabolism). It also lacks genes encoding proteins involved in pentose phosphate pathways. However, the genes for glyoxylate cycle (B9N43_03845, 03880, 03890, 05130 and 15570) and propanol-CoA metabolism (B9N43_05915, 06245, 11225, 11235 and 11240), might be able to compensate the potential deficiency of oxaloacetate, linking fatty acid degradation processes to part of the TCA cycle for producing ATP, FADH2 and NADH. Unlike the versatile anaerobic androgen degrader, *Comamonas testosteroni*, strain DHT3 does not have the complete genes for aerobic degradation of aromatics.

The ammonia can be acquired by the dissimilatory nitrate reduction (B9N43_01365, 01370, 07440, 07445, 07455, 14760 and 14775). For anaerobic growth, nitric oxide could be produced due to nitrate reduction and denitrification (B9N43_07235, 07440, 07445, 07455, 08060, 09155, 13500, 14760 and 14775), but gaseous nitrous oxide and nitrogen are not expected since the DHT3 genome lacks the gene of nitric oxide reductase subunit C. Molybdenum is an essential cofactor of the enzymes involving denitrification and the genes for molybdate transport system are identified (B9N43_04295 ~ 04305). The strain DHT3 is not able to use sulfate as terminal electron acceptor due to the lacking of anaerobic sulfur reduction and sulfate transport system in the genome.

Vitamins are essential biomolecules required for cellular metabolism. Among which, cobamides such as cobalamin are involved in biosynthesis of methionine and fatty acids in organisms among all the three domains of life (Fang *et al*., 2017). Although strain DHT3 possesses complete genes in the lower pathway for cobamide assembly from cobyric acid, the aliphatic side-chain bridge, and lower axial ligand (B9N43_07700, 08460, 10030, 10270 ~ 10280, 10430, 10480, 11085, 11090, 11100, 14110 ~ 14120 and 16605), strain DHT3 genome lacks many genes for *de novo* biosynthesis of cobyric acid via either the aerobic biosynthetic pathway or the anaerobic biosynthetic pathway. Nevertheless, complete set of biosynthetic genes for biotin (from pimeloyl-CoA to biotin; B9N43_03065, 03075, 03090 and 10790), pantothenate (B9N43_04665, 04670, 07415, 07420, 07425 and 08335), and p-aminobenzoic acid (B9N43_04985, 06085, 07405, 07565, 11365, 13845 and 14435) were identified.

### Genome of *Denitratisoma oestradiolicum* DSM 16959

After quality trimming, 13,678,478 reads were generated by the Illumina sequencing systems. The bacterial genome was assembled *de novo in silico* using ALLPATHS-LG, resulting in 65 contigs (>1,000 bp) with an N_50_ length of 170,890 bp. The genome was also annotated using the NCBI Prokaryotic Genome Annotation Pipeline and deposited as accession number of NCXS00000000. The assumed genome size of DSM 16959 is 4,144,705 bp with a G+C content of 62.0 %. Up to 3,680 protein coding genes, 5 rRNA operons (5S, 16S and 23S), 53 tRNA and 47 pseudogenes were identified. Reported as an estrogen-denitrifying bacterium (Fahrbach *et al*., 2006), several steroid degradation genes were identified in DSM 16959 genome (Table 3).

**Table 3.** Putative genes involved in the steroid degradation of three denitrifying bacteria, *Denitratisoma* sp. strain DHT3, *D. oestradiolicum* DSM 16959 and *Steroidobacter denitrificans* DSM 18526. Number in each parenthesis indicates total number of the involved genes.

| Function | Putative enzyme | Locus tag of strain DHT3 | Locus tag of DSM 16959 | Locus tag of DSM 18526 |
| --- | --- | --- | --- | --- |
| **Steroid A/B-rings degradation** | 1-testosterone hydratase/dehydrogenase | B9N43_1910~1920 (3) | CBW56_18555~18565 (3) | ACG33_RS03375, _RS03380, _RS03395 |
| | 3α-hydroxysteroid dehydrogenase | B9N43_03425 and B9N43_11350 | CBW56_10805 and CBW56_13950 | ACG33_RS09110 |
| | 3-oxosteroid $\Delta^1$-dehydrogenase | B9N43_04410 | CBW56_04345, _17020, _15965 | ACG33_RS00240, _RS03450 |
| | 3β-hydroxysteroid dehydrogenase | B9N43_16370 | CBW56_03310 | ND |
| | 3-oxosteroid $\Delta^4$-dehydrogenase | B9N43_15155 | CBW56_13900 | ND |
| **Steroid C/D-rings degradation** | Enzymes involved the HIP catabolism | B9N43_04420~04465 (10) | CBW56_04290~04335 (10) | ACG33_RS00310, _RS00315, ACG33_RS00335~RS00355 (5), ACG33_RS00370, _RS00315, ACG33_RS00745 |
| **Steroid side chain degradation** | Steroid C25 dehydrogenase | B9N43_01670~01680 (3)<br>B9N43_05455~05470 (4)<br>B9N43_11160~11175 (4)<br>B9N43_15060~15070 (3) | CBW56_04635~04645 (3)<br>CBW56_13870~13885 (4)<br>CBW56_02265~02275 (3)<br>CBW56_02870~02880 (3)<br>CBW56_17885~17895 (3) | ACG33_RS10675~10690 (4) |
| **Anaerobic estrogen catabolism** | Uncertain | B9N43_10285~10330 (10)<br>B9N43_08210~08240 (7) | CBW56_16910~16950 (9)<br>CBW56_09525~09555 (7) | ACG33_RS12610~12650 (9)<br>ACG33_RS11635~11665 (7) |

### Comparative genomic analysis

To further mine genes involved in steroid anaerobic catabolism, 6 bacterial genomes of different steroid degraders were chosen for this analysis. Up to 689 homologous gene clusters are shared among these genomes (Figure 2), including the genes involved in the 2,3-seco pathway and in steroid C/D-rings degradation. The latter implies that HIP might be the common metabolite in either aerobic or anaerobic degradation pathways. Based on their physiological characteristics and genomic difference analysis, we found that there are 45 homologous gene clusters were only identified in these estrogen denitrifying bacteria (strain DSM 18526, DSM 16959 and DHT3), and 41 of them are single copy genes. Strikingly, 10 of these shared genes are located in two confined area in each of the genome, and most of their function are unknown through this analysis (Table S1; cluster_name: cluster0020 ~ 0028). As a result, these 10 genes might be the key to unveiling the biochemical pathway of anaerobic estrogen degradation (Table S1 and Table 3).

**Figure 2.**
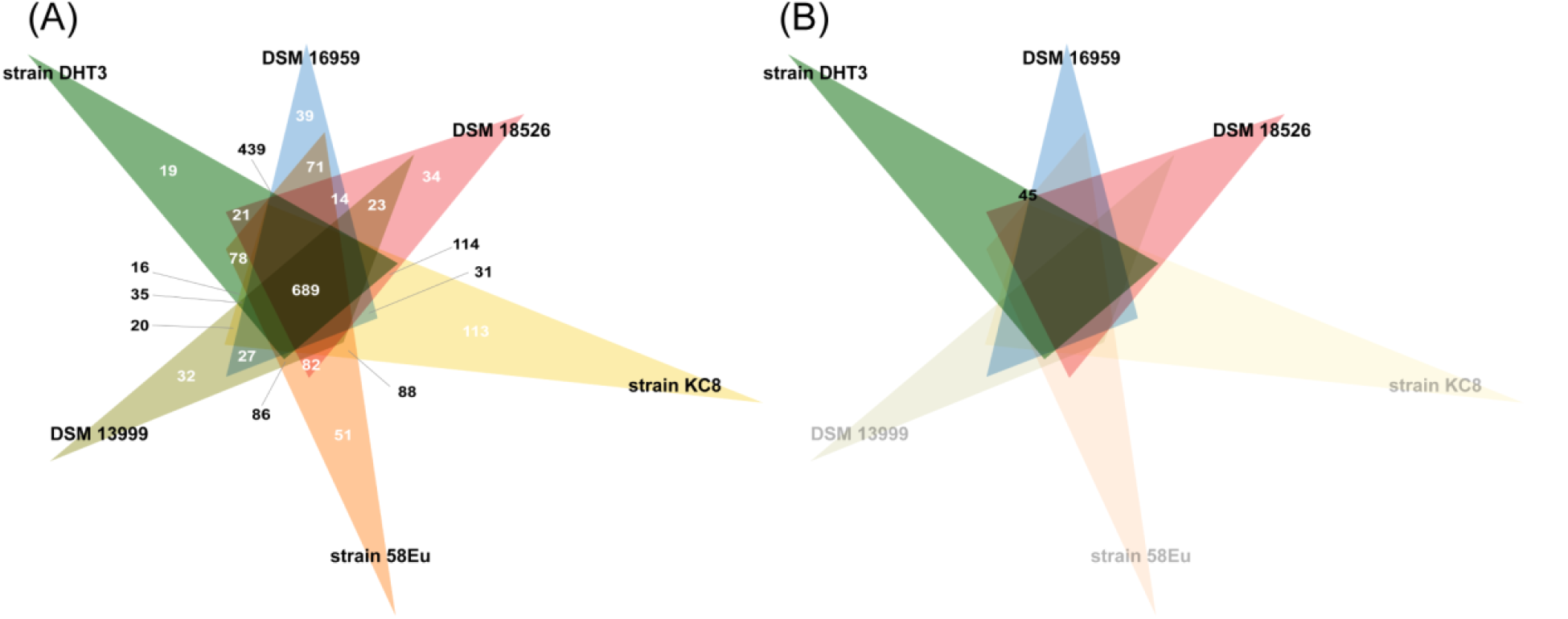
Venn diagram of shared orthologus clusters (A) among the six aerobic and anaerobic steroid degraders, and (B) among three estrogen-degrading denitrifies.

